# Autonomous control of extrusion bioprinting using convolutional neural networks

**DOI:** 10.1101/2024.12.07.627315

**Authors:** Daniel Kelly, Vasileios Sergis, Laura Ventura i Blanco, Karl Mason, Andrew C. Daly

**Affiliations:** Biomedical Engineering, School of Engineering, College of Science and Engineering, University of Galway, Galway, Ireland; CÚRAM, Research Ireland Centre for Medical Devices, University of Galway, Galway, Ireland; School of Computer Science, College of Science and Engineering, University of Galway, Galway, Ireland

**Keywords:** 3D bioprinting, closed-loop bioprinting, convolutional neural networks, machine learning, additive manufacturing, quality control

## Abstract

Extrusion bioprinting technology suffers from reproducibility challenges due to the open-loop nature of current hardware systems. Here, we present a novel AI-powered extrusion bioprinting platform with integrated real-time quality monitoring and automated error correction capabilities. To achieve this, we engineered a custom bioprinting system with an integrated camera for continuous process monitoring and trained convolutional neural networks (CNNs) to classify the extrusion process in real-time. The CNN models, including Xception and ResNet, were trained on a combination of real and synthetic data to classify extrusion quality (good, over, or under) across various printing scenarios, including single-line and infill patterns. Notably, transfer learning, utilizing synthetic data for initial training followed by refinement with real-world data enhanced classification accuracy, with the Xception model displaying 90% accuracy for single-line extrusion and 75% for infill extrusion. This intelligent monitoring system was then coupled with a closed-loop control system that dynamically adjusted extrusion parameters on-the-fly to correct errors. The platform successfully corrected both over- and under-extrusion errors for alginate and collagen bioinks with varying rheological properties, demonstrating adaptability to unseen materials. Importantly, extrusion errors were corrected within ∼10 seconds. This novel closed-loop bioprinting platform represents a significant advance over traditional open-loop systems.

## 1. Introduction

Bioprinting technology holds tremendous potential for developing artificial tissues and organs that mimic the complexity of their native counterparts. Through the layering of cell-laden bioinks using extrusion or lithography technology, bioprinting enables spatial patterning of multiple cell populations in prescribed configurations. Recent advances have enabled the bioprinting of anatomically accurate constructs mimicking the geometry of an entire heart [1,2]. However, despite these notable advances, translating this promise into robust and reproducible bioprinting protocols for organ manufacturing presents significant challenges [3,4]. A central challenge lies in optimising bioprinting protocols and bioink formulations to enable manufacturing of anatomically accurate structures, while supporting high cell viability and post-printing tissue maturation [5,6]. This is complex due to the multitude of interdependent parameters that need to be optimised for a given bioink formulation (e.g., flow rate, print speed, print height, viscosity, temperature, layer height), necessitating extensive experimentation by specialists [7,8]. Even when optimal processing conditions have been identified, bioink inconsistencies or environmental changes can introduce extrusion errors, leading to bioprinting failures. These challenges are exacerbated by the high cost of bioink materials and the stringent regulatory requirements that will exist for bioprinted Advanced Therapy Medicinal Products (ATMPs), underscoring the need for reproducible and reliable protocols. The limitations described above largely stem from the open-loop nature of existing bioprinting hardware, which lacks real-time process monitoring and adaptive control over extrusion parameters. Process parameters are fixed before printing, which prevents live editing of extrusion parameters to correct errors. Addressing this open-loop limitation will be crucial for advancing bioprinting beyond academic labs and into practical, industrial applications.

The challenges with open-loop printing methods extend beyond 3D bioprinting to the general additive manufacturing industry, regardless of the inks utilised or the application [8–14]. Recent attention has been directed towards implementing monitoring and closed-loop control technology to improve the quality of 3D-printed components. In contrast to traditional open-loop 3D printing, where parameters are fixed before printing, closed-loop 3D printing refers to techniques that dynamically adjust printing parameters to compensate for changes such as printing defects, ink flow irregularities, nozzle functionality, and spatial displacement errors [9]. In the fused-deposition modelling field, closed-loop control mechanisms have been implemented using traditional [15–19] or AI-based [20–24] approaches, incorporating sensors into the printing process to evaluate the quality of the print [10,25]. Camera-based sensors are the most widely adopted, with some examples of combining camera feeds with convolutional neural networks (CNNs), a powerful approach to identify printing defects and provide feedback to the material-feeding and motion-control systems to correct printing errors [16,20,26–30]. More recently, closed-loop extrusion controllers applied through reinforced learning and trained on simulation data have been developed [31]. Although closed-loop techniques are becoming more widespread in the general additive manufacturing sector, there has been limited adoption for bioprinting hardware. While early work has shown the potential of camera systems and machine learning for monitoring extrusion bioprinting and identifying anomalies [32,33], current hardware platforms often rely on cameras positioned at oblique angles and large distances from the printing nozzle. This limits high-resolution monitoring and restricts error classification to post-printing analysis or after completing entire layers within the print. Crucially, this limits the potential for on-the-fly error detection and extrusion rate adjustments during the bioprinting process.

Here, we developed a novel bioprinting platform with integrated computer vision and machine learning algorithms that can autonomously detect and correct extrusion errors in real-time. This was achieved by developing novel hardware with an integrated camera for real-time continuous monitoring of the extrusion process. Using a combination of experimental and synthetic data, we then trained CNNs that could rapidly identify and classify extrusion errors and inconsistencies in filament diameter, including under-extrusions and over-extrusions, based on real-time camera feeds. Building on these error detection capabilities, we then developed closed-loop extrusion controllers that dynamically adjust flow rate to correct the identified errors in real-time. We validated the platform’s ability to detect and correct extrusion errors across a range of bioink formulations with diverse rheological properties, demonstrating its potential to enhance the reliability and reproducibility of bioprinting with challenging bioink formulations. This fully automated, closed-loop platform represents a significant advance beyond the state-of-the-art for the bioprinting field, which to date has relied on open-loop hardware.

## 2. Results and Discussion

### 2.1 Training convolutional neural networks for extrusion error detection during bioprinting

We first focused on training CNN models that could rapidly identify extrusion anomalies during bioprinting. To enable this, we leveraged a bioprinting hardware platform with integrated quality monitoring functionality that was recently developed in our group [34]. This hardware solution comprises a readily available and low-cost bed-slinger fused deposition modelling 3D printer (Prusa MK3S+) with the thermoplastic extruder swapped for a progressive cavity pump extruder (Puredyne kit B), thus converting the printer into a bioprinter (Figure 1a). The printer is further modified with a second x-axis located below the extruder x-axis that is fitted with a Raspberry Pi 1.6-megapixel global shutter camera and macro scale lens that focuses directly up towards the tip of the extruder and follows its movements. The result is a system that can print on any transparent medium with an unobstructed view up at the extruder depositing material (Figure 1a, Movie S1). For initial testing, a 100 mg/ml alginate bioink was employed, and red dye was added for visualisation. This is a unique perspective not seen in prior bioprinting research, where close-view imagery is generally taken at an angle from above or perpendicular to the extruder, often at larger distances that prevent high-fidelity extrusion monitoring [32,33]. Leveraging this platform, we focused on training CNNs that could detect extrusion errors from the live imaging feed. We focused on developing CNNs that could classify the current state of the extrusion process as good, over, and under, with good extrusion defined as bioink deposition where the filament width matched the bore of the extruder nozzle. All experiments were performed with a 20G nozzle (603𝜇𝑚 bore diameter), but due to the nature of our implementation, the platform will automatically adjust the definition of good extrusion depending on the bore diameter of the nozzle employed. To effectively train a CNN, image data is required to converge upon the optimal model parameters such that images can be accurately classified from their features. We created two large, labelled datasets of images from the vantage point of our hardware system. The first dataset comprised real-world images of extrusion outcomes taken directly from the computer vision system (Figure 1a). However, generating real-world data with sufficient diversity of training examples for CNN model training is costly in terms of materials and time. To alleviate this burden, we developed a 3D simulation of the bioprinting process where material parameters can be augmented at will, prints can run faster than they would in real life, and each image produced can be precisely labelled as good, over or under as the extrusion quality is always known (Figure 1b). This simulation was built in Blender [35] and uses the in-built fluid modelling package to approximate different extrusion bioprinting outcomes (Figure 1a, b).

**Figure 1:**
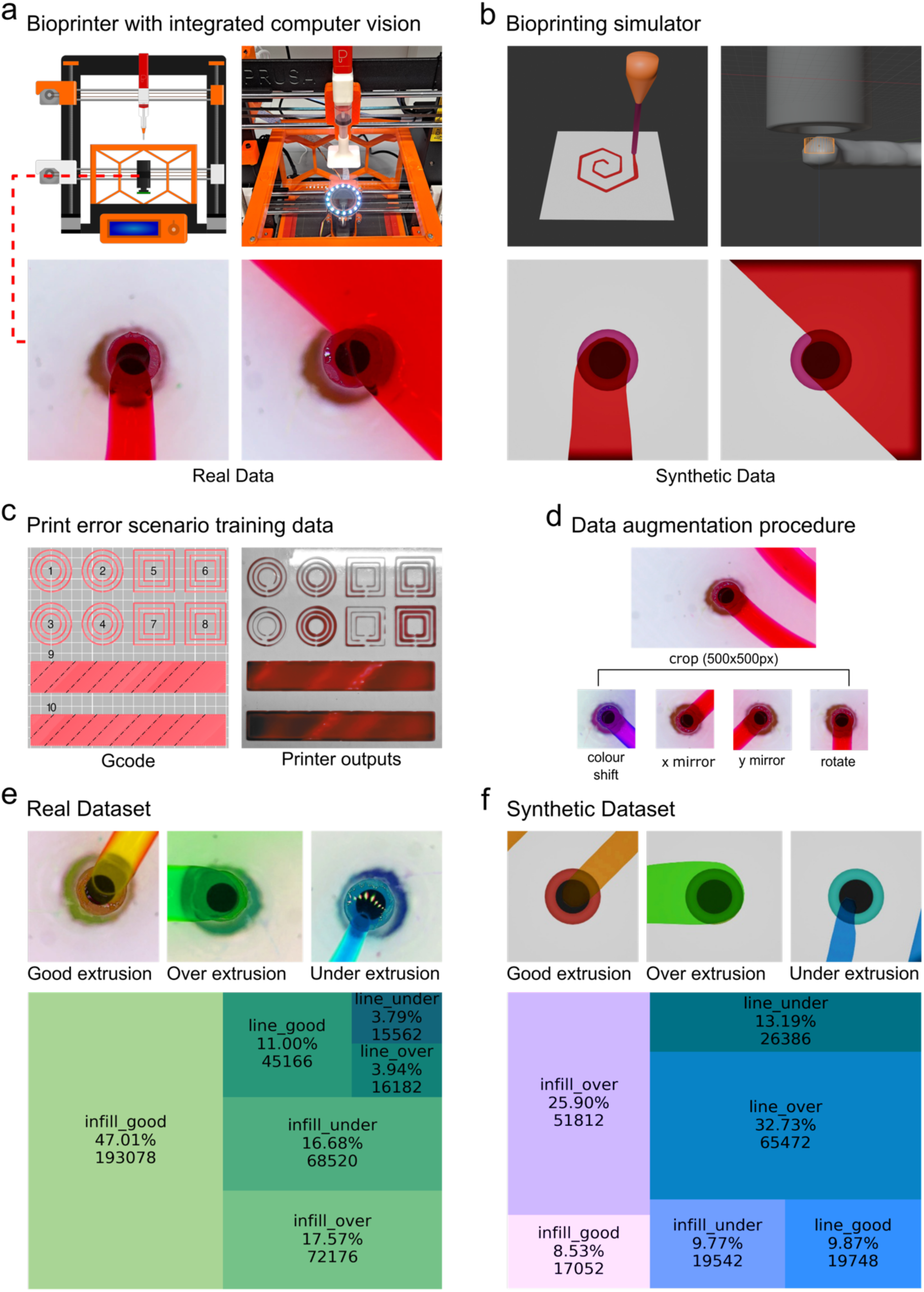
Data generation process for training CNNs: **(a)** (top) Schematic and picture of developed bioprinting hardware showcasing the cloned X axis with a camera, transparent build plate, and Puredyne extruder. The extrusion nozzle was 20G (603 𝜇𝑚 bore diameter). (bottom) Example images of extrusion from the macro zoom camera. **(b)** (top) Recreation of bioink extrusion through the bioprinting nozzle in Blender using built-in Fluid Implicit Particle (FLIP) fluid modelling. (bottom) Example images showing the high resemblance between the real and synthetic imagery. **(c)** (left) Visualisation of the g-code used to gather labelled experimental data. Shapes 1 to 8 are split in half with altered extrusion rates alongside normal extrusion. Shapes 9 and 10 have altered extrusions in each segment marked by dashed lines. (right) resulting prints that were created using these g-code files with the dyed 100 mg/ml alginate bioink. **(d)** Examples of the image augmentation procedures that were applied to both datasets to increase the diversity of the training data. **(e-f)** Resulting images that were created after augmentation of the real and synthetic datasets, respectively, and the ratio and image count of each class contained in the datasets.

To accurately label the real-world images from the bioprinting hardware, specific patterns were printed with regions of poisoned g-code, that is, areas where the extrusion value was purposely set too high or too low (Figure 1c). The images gathered from those regions were labelled in correspondence to the type of poisoning. The real-world data was produced using concentric circle geometries where each half of the circle contained good extrusion, and the other half contained lightly under, lightly over, very under, and very over extrusion (Figure 1c, 1-4). This process was repeated for square patterns (Figure 1c, 5-8). These print geometries represented single filament extrusion outcomes that generally only represent borders of a structure, so to further augment the dataset and ensure it was representative of real-world bioprinting scenarios, we included solid structure prints with infill patterns (Figure 1c, 9-10). Within these prints, the dashed lines represent areas of alternating good, over, or under extrusion (Figure 1c, 9-10). All images in both datasets were further processed using data augmentation techniques to increase the data yield and variability. The augmentations performed included cropping, colour shifting, mirroring in the x and y axis, and rotation (Figure 1d). This resulted in synthetic and experimental datasets consisting of two extrusion scenarios (single line or infill extrusion) that were then further categorised as good, over or under extrusion (six categories total) (Figure 1e, f). The resulting datasets have been made openly available [36]. The real dataset images were created using the alginate bioink containing red dye.

The CNNs were created using Python and the Tensorflow [37] machine learning library. Four high-performing architectures from the ImageNet Large Scale Visual Recognition Challenge [38] (ILSVRC) were chosen to be trained on our datasets: Xception [39], ResNet50V2 [40], MobileNetV3 [41], and VGG19 [42]. These architectures come pre-built in Tensorflow for ease of training on custom datasets. The data was split 80/20 into a training dataset and a hold-out dataset for testing after training. The training dataset was further split 80/20 into training images and validation images during training. The training process was configured to run for 100 epochs or until no new maximum accuracy was achieved after 20 epochs. We evaluated the accuracy, precision, recall and F1-score of each model on the hold-out test data across all permutations of infill or line extrusion on the synthetic and real datasets (Figure 2a). Additionally, we include a set of transfer-learned models that are synthetic models further trained on real data (Figure 2a). All CNN models performed well across all metrics, scoring above 90% except for synthetic infill (above 85%) and transfer VGG19 infill, which did not successfully train (Figure 2a). To further evaluate the accuracy for each of the CNN models, confusion matrices were used to compare the predicted classification to the true classification of 512 images (Figure 2b). These matrices highlight any tendencies for a model to mis-classify one class as another. For example, ResNet50V2 synthetic infill had a tendency to misclassify good extrusion as under extrusion or, to a lesser extent, over extrusion (Figure 2b). The results show a strong classification performance, with most classifications being correct. In order to understand what features have the strongest impact on a CNN’s classification decision, saliency maps [43] were produced for each model on an example of infill and line extrusion (Figure 2c). These maps highlight the attention of a model on areas of an image (most apparent for VGG19). These maps do not reflect on the model’s performance but rather serve as a means of interpreting their inner workings, which is a goal of explainable artificial intelligence (XAI). From our maps, we could interpret a strong focus on the centre of the image around the border of the extruded bioink and the area immediately around it. Having demonstrated that the CNN models could accurately classify different bioprinting extrusion scenarios on previously unseen hold-out labelled data, we next sought to evaluate performance on real-world bioprinting hardware.

**Figure 2:**
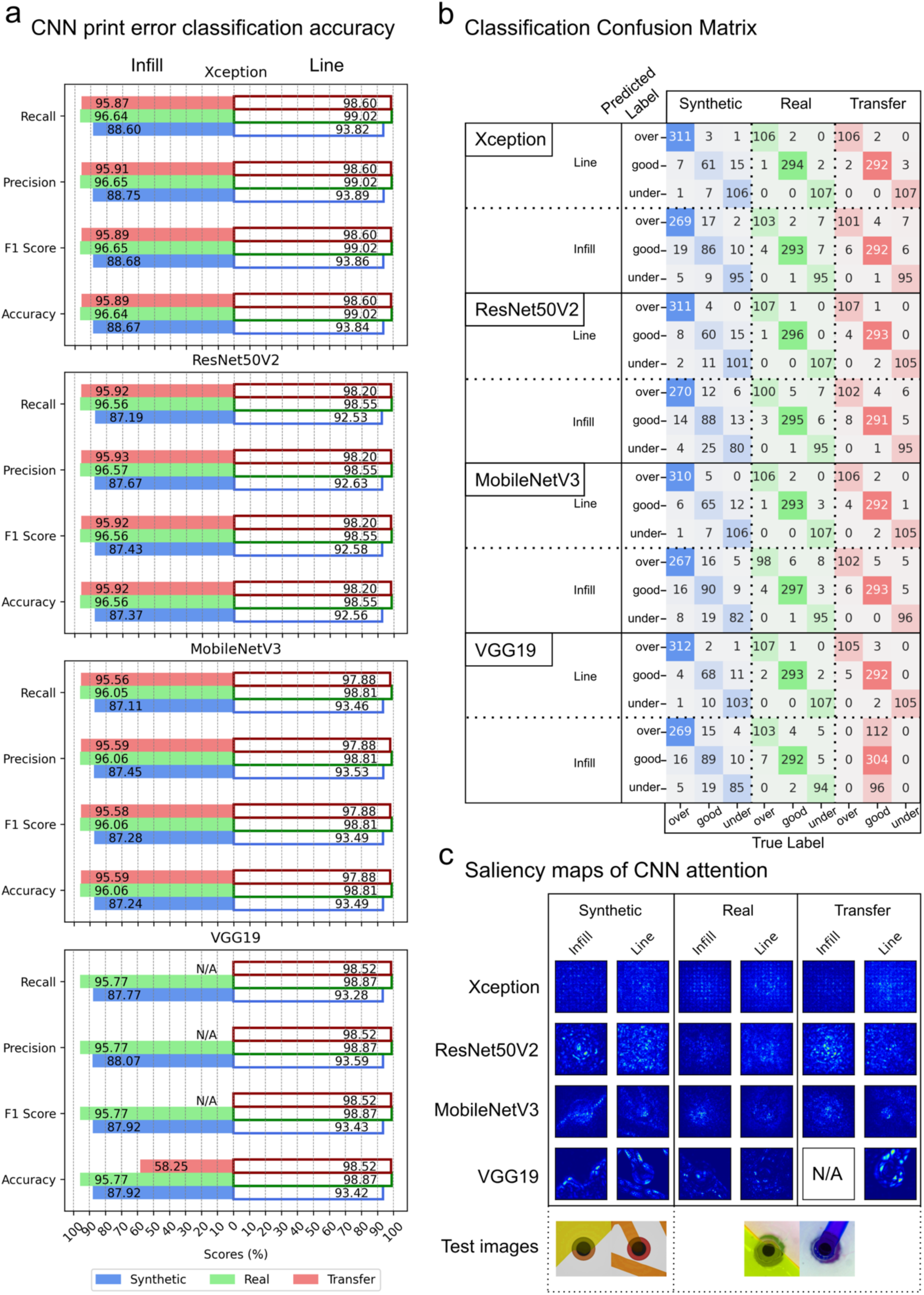
Evaluation of four CNN models trained on synthetic data only, real data only, and synthetic followed by real data (transfer learning): **(a)** Evaluation results of all models on holdout datasets, reporting accuracy, F1, precision and recall. Blue bars are models trained on synthetic data only, green bars are real data only, and red bars are trained on synthetic data first, then the real data (transfer learning). Each of the four architectures are split into the infill and line models and placed beside each other on a diverging x-axis. VGG19 infill transfer learning model is not displayed as it failed to train **(b)** Confusion matrixes for each model, showing the true classification for each predicted classification. The models are sorted with the same colour coding and split for infill and line prints. **(c)** Saliency maps of the attentions of each model. VGG19 infill transfer learning model is not displayed as it failed to train. The images used to produce the saliency maps are placed under the corresponding maps.

### 2.2 Deployment of trained convolutional neural networks on bioprinting hardware for real-time error detection

Next, we evaluated CNN model accuracy on the physical bioprinting hardware setup with integrated computer vision (Figure 3a). Live imagery from the computer vision camera is more variable than the hold-out dataset as there are changes in lighting, unseen printing paths will be used, and minor visual artefacts can arise, such as dust and liquid droplets. The validation hardware comprises the modified printer previously described in section 2.1(Figure 3a ii and iii), a separate, more powerful computer (Macbook M1) running the Tensorflow backend for the CNN classification models (Figure 3a v), and a monitor for live analysis of the print by the operator (Figure 3a i). The Raspberry Pi sends a vector representation of the Region of Interest (ROI) from the captured image over a User Datagram Protocol (UDP) packet to the computer, which is running a server awaiting these small images (Figure 3a v). The computer processes the image to undergo classification by the model running in Tensorflow, and a response is sent back to the Raspberry Pi containing the predicted label and confidence values of that prediction. These values are displayed on the monitor via a Heads Up Display (HUD) over the live feed of the print, where the operator can monitor the current extrusion classification and a percentage breakdown of each classification across the entire print (Figure 3a iv).

**Figure 3:**
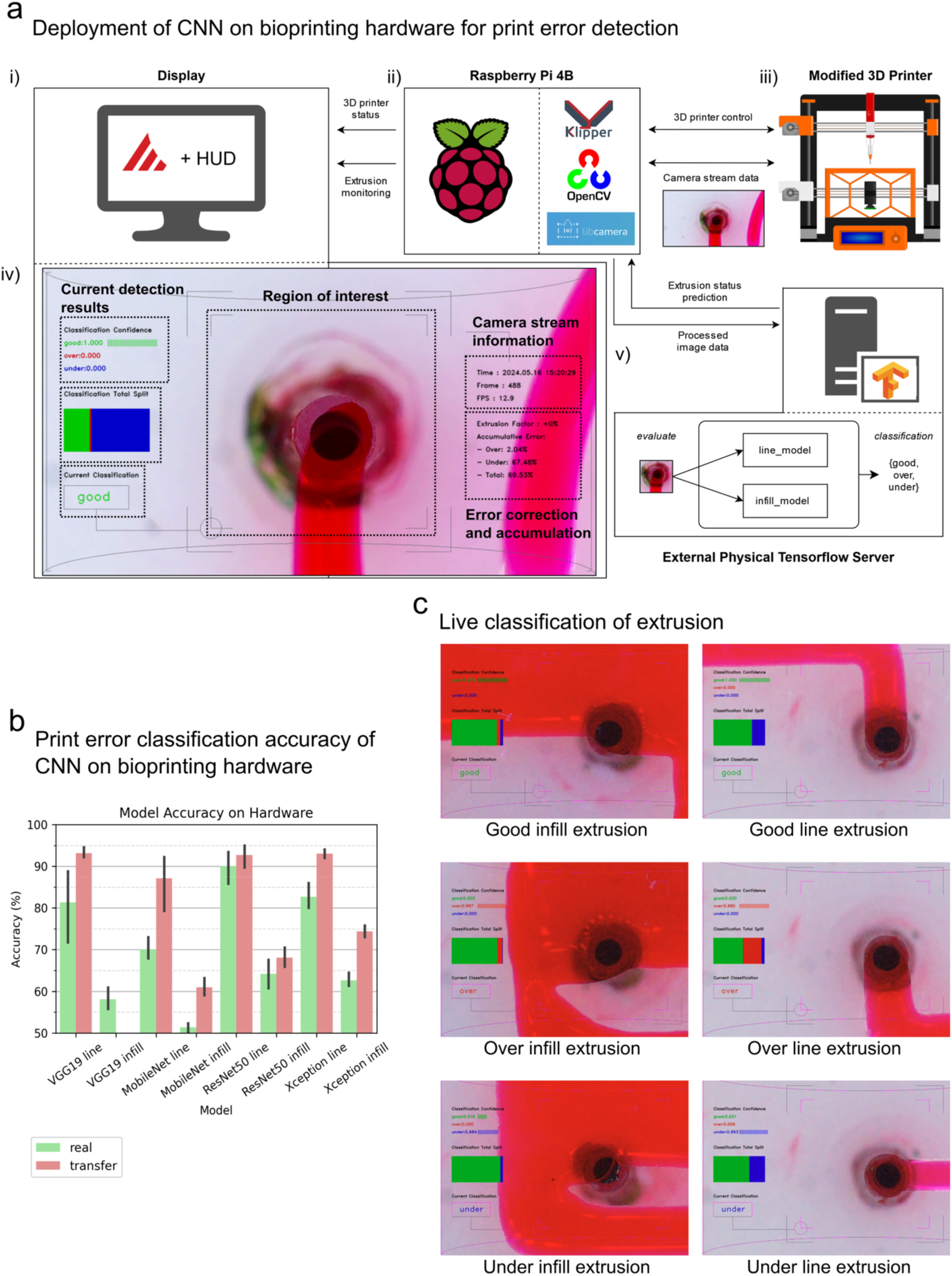
Validation of trained CNN on live printing process: **(a)** System overview of experimental setup. (i) User interface running Mainsail (Klipper graphical user interface) and our extrusion monitoring HUD. (ii) Raspberry Pi 4b handling movement and vision tasks using Klipper, OpenCV and libcamera. (iii) modified bioprinting hardware. (iv) Extrusion monitoring HUD with panels showing the current detection results (classification confidence, ratio of classifications over the print, and classification label), the camera information (time, current frame number, and frames per second), and the accumulated error over time. (v) Tensorflow server that runs on a separate computer for classification of images sent from printer. The server parses the image to the appropriate model for classification before sending a response back to the printer. **(b)** Accuracy of the detection system on prints with known areas of bad extrusion. Each model performed five runs of the print. Error bars represent 95% confidence interval. The experiment was performed using a 20G extrusion nozzle (603 𝜇𝑚 bore diameter). **(c)** Examples of correct classification of extrusion types on each type of print by the best-performing architecture (Xception).

The accuracy of the CNN models when deployed on the live extrusion feed was measured using a similar procedure to how the images for training were gathered. The printer executed simple print paths with known poisoned areas, and every detection from the CNN model was logged. The logged detections were then checked against the known extrusion value based on location within the path, and this data was used to compare the print extrusion error detection accuracy for each CNN model (Figure 3b). Five runs were completed for each model. It should be noted that this method does not account for correctly classified extrusion in an area designated with another label, e.g. when printing in a good extrusion zone, there may be natural occurrences of over or under-extrusion that get classified correctly, but because they are in the good extrusion zone, this is counted as a negative result. This method of evaluating the CNNs was chosen to streamline the validation process over the alternative, which would be to manually check hundreds of classifications. Interestingly, there was a clear trend of the transfer-learned CNN models performing better than those only trained on the real dataset (Figure 3b). When comparing the different CNN models, Xception and ResNet50 demonstrated the highest classification accuracy, with accuracies above 80% for single-line extrusion scenarios. (Figure 3b). Higher classification accuracies were observed for the line geometries compared to infill (Figure 3b). For example, the accuracy of the transferred trained Xception model was 90% for single-line extrusion and 75% for infill extrusion conditions (Figure 3b). This is likely due to the more complex and variable extrusion imagery produced during infill extrusion, where the newly deposited bioink fuses with previously deposited ink. Altogether, our results demonstrate that our trained CNN models could be deployed to detect extrusion errors during bioprinting, with the transfer learning trained ResNet50 and Xception models being the most promising. Building on this, we next sought to use our computer vision platform and error detection algorithms to introduce extrusion corrections during bioprinting.

### 2.3 Closed-loop bioprinting error detection and correction using convolutional neural networks

Next, we developed an extrusion error correction system using the classification output of the best-performing CNN models (Xception and ResNet50) (Figure 4a). This algorithm involved classifying the state of the extrusion process as either good or over/under extrusion, and when in a state of over/under extrusion, implementing extrusion adjustments until the extrusion state returned to good (Figure 4a i). For initial testing of this detection and correction model, simple rectangle geometries were printed, and the g-code was poisoned at specific locations to introduce both over- and under-extrusion errors. The print path was split into 1mm sections such that the printer would extrude bioink for a linear distance of 1mm, send an image of the extrusion outputs to the Tensorflow server, and then receive a classification response before printing the next 1mm (Figure 4a ii). The round-trip time for this process was tuned to 1 second for fast print error detection and smooth extrusion adjustments. The number of extrusion observations along the print path could be increased depending on the nature of extrusion errors for a specific bioprinting application. As the system sends g-code line-by-line in a just-in-time fashion, the correction operation can directly alter the extrusion value of the following line of g-code to ensure immediate response. A global extrusion factor is increased when the classification is currently under and decreased when over. The extrusion value is multiplied by this factor to adjust the amount of material deposited. The extrusion factor is incremented in steps of 15%, starting at 100% every time a detection is made, with upper and lower limits of 20% and 200% for the final factor to protect against over-correction. This process is faster than using the in-built extrusion flow percentage adjustment in Klipper (g-code command M221) as it edits the g-code before being processed by the printer.

**Figure 4:**
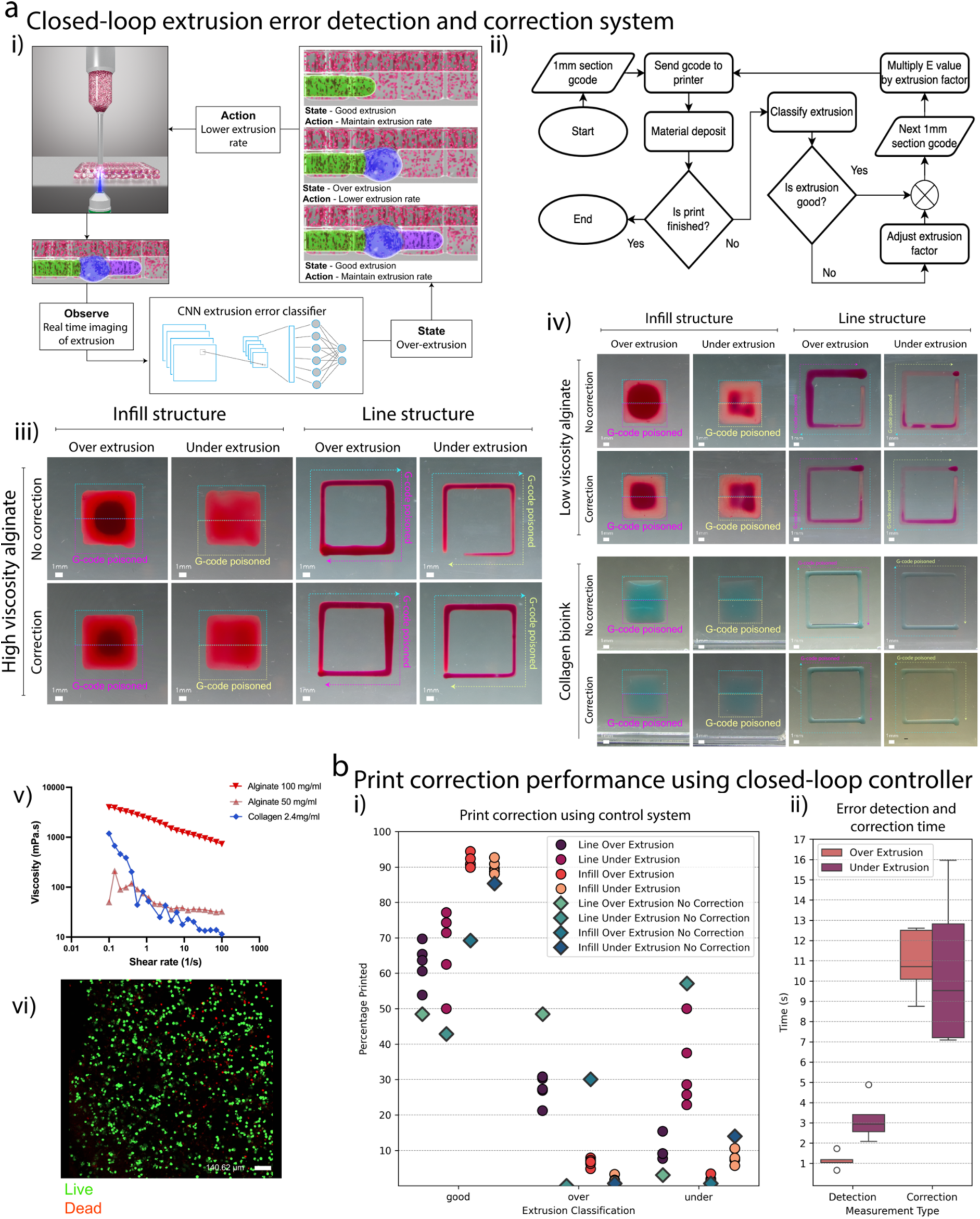
Correction and Robustness testing of CNN on materials unseen in the training process: **(a)**(i) Desired process of controlling extrusion using computer vision where the system observes the material deposition, declares the current state of extrusion, and performs some action to alter the extrusion. (ii) Specific control process applied to our experimental setup to correct extrusion errors in real-time. (iii) Resulting prints of infill and line with high viscosity alginate (100 mg/ml) undergoing correction. The top row receives no correction from the system. The over and under extrusion starts halfway on each print. Scale bars represent 1mm. (iv) Robustness test on unseen materials; low viscosity alginate (50 mg/ml) and collagen bioink (2.4 mg/ml). Scale bars represent 1mm. (v) Rheological characteristics of each bioink demonstrated using a shear rate sweep. (vi) Live/dead staining of cells (human induced pluripotent stem cells; 5 million/ml) within the collagen bioink post-bioprinting for an infill geometry. Scale bar represents 140 μm. All validation prints were performed using a 20G extrusion nozzle (603 𝜇𝑚 bore diameter). **(b)**(i) Effect of control system on print accuracy for high viscosity alginate. The diamond-shaped datapoint in each column represents the overall accuracy of the print with no active corrections. The accuracy is determined by the CNN detection system results at the end of print for five runs. The circular data points indicate overall print accuracy when the closed-loop controller (detection and correction) is turned on. (ii) Time measured to detect and correct over and under extrusion over the five runs of single-line extrusion.

When deployed on poisoned g-code files, we found that the closed-loop controller could correct extrusion errors and improve overall print quality (Figure 4a iii, Movie S1). Compared to standard open-loop bioprinting (no correction), our closed-loop controller enhanced extrusion quality across both line and infill structures printed using the dyed alginate bioink (Figure 4a iii, 4b i). For example, for line structures containing a deliberate over-extrusion, the closed-loop controllers increased the percentage of the print classified as good from 50% up to 65% (Figure 4b i). Similarly, for infill structures containing a deliberate over-extrusion, the closed-loop controller increased the percentage of the print classified as good from 70% up to 90% (Figure 4b i). The closed-loop controllers also improved overall print quality for infill and line structures containing deliberate under-extrusions (Figure 4b i). It took an average of 1 and 3 seconds to detect erroneous extrusion from the known point of poisoning for over and under-extrusion, respectively (Figure 4b ii). This translates to 1 and 3 cycles of detection to identify the error. Correction of the extrusion errors with the closed-loop controller took 10.75 and 9.5 seconds for over and under-extrusion, respectively (Figure 4b ii). Future work will focus on optimising the control scheme and communication time between the different modules to speed up the controller’s response time. Despite the computational lag, the closed-loop controller substantially increased print quality for both single-line and infill structures with over and under-extrusion errors.

The initial validation tests were performed using the 100mg/ml alginate bioink containing red dye, the same material used in the CNN training data. To evaluate the adaptability of our closed-loop controller to other bioink formulations, the live correction tests were first repeated using an unseen alginate bioink with a lower viscosity (50 mg/ml) (Figure 4a iv, v, Movie S2). We also evaluated the controller’s performance on an unseen collagen bioink containing a blue dye (2.4 mg/ml) (Figure 4a iv, v, Movie S3). We found that the error detection system could successfully classify the extrusion outputs for these unseen bioinks (Movie S2 and S3). We also found that the closed-loop controller improved print quality for both unseen bioink formulations, with improvements in print quality for infill and line structures containing over and under-extrusion errors (Figure 4a iv, Movie S2). It was observed that the less viscous bioinks were not as responsive to the correction inputs, and in the case of the collagen, the poisoning (Figure 4a iv, Movie S3). However, improvements in print quality were clearly observed for the collagen bioink in under-extrusion conditions (Figure 4a iv). It should also be noted that the CNN error detection system could respond to the collagen bioink containing a blue dye. This result was expected due to the data augmentation techniques employed during CNN model training (Figure 1d). The blue dye (Brilliant Blue FCF) is compatible with cells and did not negatively influence the viability of encapsulated human induced pluripotent stem cells (5 million/ml) during extrusion with the collagen bioink, confirming compatibility with bioprinting applications (Figure 4b vi). Together, these results demonstrate that the controller can adapt to unseen bioinks with varied rheological properties, confirming the robustness of the closed-loop detection and correction controller.

This work advances the state-of-the-art in extrusion bioprinting by introducing closed-loop process controls that can detect and correct extrusion errors using CNNs. While machine learning has been previously employed for anomaly detection and quality monitoring in bioprinting [32,33], existing approaches have focused on assessing quality across whole print layers using wide camera views, limiting the potential for real-time error correction within individual filaments. Our high-resolution quality monitoring platform overcame this limitation, enabling on-the-fly extrusion error detection and correction. With error detection and correction cycles occurring within 8-12 seconds, depending on the nature of the initial extrusion error, our system offers a significant advantage over approaches that rely on post-print or layer-by-layer analysis, enabling rapid adjustments and preventing the propagation of errors throughout the fabrication process. This real-time error correction capability is important in bioprinting applications, where the high cost of bioink materials necessitates precise control and minimisation of waste.

The ability of our system to operate on unseen bioinks is an important step towards more versatile and adaptable bioprinting technology that can autonomously adapt process parameters for new bioinks. This capability stems from our use of a transfer learning approach on augmented datasets of both synthetic and real images, which allowed the CNN models to extract relevant features and adapt to variations in bioink composition (rheological properties, colour) without requiring extensive retraining for each new material. The model could be further enhanced beyond the results presented in this work by continuous transfer learning on a wider dataset of different materials. Beyond enabling real-time extrusion error detection and correction, these capabilities could automate the parameter optimisation process for new bioinks and/or bioprinting protocols. Such an automated approach to parameter optimisation could accelerate the development of new bioprinting protocols, reducing the reliance on manual tuning and facilitating the exploration of a wider range of bioink formulations and printing parameters.

A current limitation of our work is that we have only demonstrated extrusion detection and correction for single-layer prints. This is because our current camera system captures high-resolution 2D representations of the extrusion process, which was an intentional choice to investigate the highest fidelity viewpoint of bioink extrusion. While this viewpoint is highly beneficial for first-layer monitoring and fine-tuning extrusion parameters, it ultimately cannot perform the same operations on subsequent printing layers within a 3D construct. Future work will explore integrating multiple camera angles and volumetric microscopy techniques into our platform.

## 3. Conclusion

We have developed an advanced closed-loop bioprinting platform powered by CNNs capable of autonomously detecting and correcting extrusion errors in real-time. By training our CNN models via transfer learning using a combination of real and synthetic data, it was possible to achieve high error classification accuracies for both single filament and infill extrusion scenarios. Crucially, the platform was capable of optimising and correcting extrusion outcomes for unseen bioink formulations with varied rheological properties, highlighting adaptability to diverse materials and demands. This intelligent system represents a significant advancement over conventional open-loop bioprinting methods, which lack real-time feedback and error correction capabilities. This technology holds great potential for improving the accuracy, reliability, and reproducibility of extrusion bioprinting processes, a key consideration for industrial and GMP-compliant manufacturing.

## 4. Experimental section

### 3D bioprinting setup

The hardware used for generation of images for CNN training and subsequent bioprinting with live error correction is based on a Prusa MK3S+ FDM 3D printer (Prusa3D). This printer uses a cartesian kinematic system with independent control of the X, Y and Z axis where the extruder is attached to the X and Z axis and the build surface is attached to the Y axis, commonly referred to as a bed-slinger. To achieve a view of the extrusion from directly under the extrusion nozzle, a second X-axis was installed that follows the movement of the primary X-axis. On this cloned axis, a Raspberry Pi global shutter camera (SC0715 Pimoroni) with macro lens (CAM011 Pimoroni) attached. Between the extruder and camera, a glass plate is held by a cage attached to the Y axis to act as the new printing surface. The printer is configured with Klipper for control of its movement. A Raspberry Pi 4b (4255998 Farnell) is connected in parallel with the printer mainboard for interface with Klipper and control of the camera. Raspberry Pi OS 64 bit Bookworm is installed on the Raspberry Pi and Klipper running as a background process to allow access to a standard Linux interface for use of python scripts for the control of the camera and monitoring system. The FDM extruder is replaced with a Puredyne kit B (Puredyne) LDM extruder.

### Image Dataset Generation

Images of real prints were taken using the hardware setup previously described. The camera captures images at 24 frames per second in high-definition quality (1280 × 720 px). The prints performed for dataset generation follow a specific pattern such that large groups of images can be labelled at once based on the region of the pattern they follow. These regions include extrusion purposely tampered with to produce over and under-extrusion. The bioink printed was 100 mg/ml alginate with red dye, where 500 mg of alginic acid and 50 mg of Direct Red dye were added to 5 ml of water. These labelled images are further processed by cropping around the centre of the extruder nozzle to a size of 224 × 224 px and performing one or multiple of the following image augmentation techniques: rotation, flipping, and colour shifting. The full dataset can be found here https://doi.org/10.5281/zenodo.13939763.

### Blender extrusion simulator

A simulation of the printing output was created in Blender for the creation of synthetic imagery. The simulator moves a nozzle along a defined path over a transparent surface. At the tip of the nozzle a fluid emitter spawns simulated liquid using FLIP configured to resemble the gel used in the real images. The emitter is scaled to achieve different extrusion qualities and labelled based on the scale value. The same image augmentation is performed on the images captured from the simulator.

### CNN model training

Tensorflow is used for processing of the datasets and training of the CNNs. Four CNN predefined architectures were chosen for training: Xception, ResNet50V2, MobileNetV3 and VGG19 from the built in Keras models using a Stochastic Gradient Descent optimiser. The data is split 80/20 into training and testing datasets. The training dataset was further split 80/20 into images for training and images for validating each training epoch. Each model was trained for a maximum of 100 epochs with a patience of 20 epochs for a new max accuracy value. The final models were then tested on the hold out test dataset for final accuracy, precision, recall, and F1 scores. Full details of the code can be found at https://doi.org/10.5281/zenodo.13981630.

### Human induced pluripotent stem cell expansion and maintenance

Human induced pluripotent stem cells (hiPSC) (A18945, Fisher Scientific) were seeded on Corning Matrigel (354277, Fisher Scientific) coated six-well plates. The Matrigel coating solution was prepared by adding 20 μl of Matrigel to 1ml of Gibco DMEM F-12 GlutaMAX supplement medium (10565018, Fisher Scientific). The plates were coated for 1h at room temperature and incubated at 37°C for 15min. Passage 6 (P6) hiPSCs were seeded on the plates and maintained in Gibco Essential 8 Flex Medium (A28583-01, Fisher Scientific). Media was changed daily (every other day for weekends) until the cells reached 70-80% confluency. The cells were detached using Accutase (07922, STEMCELL Technologies) through incubation at 37°C for 5mins, followed by centrifugation at 200G for 3 min. The cell pellet was then gently dissociated to prepare the cell suspension medium for the bioink.

### Bioink preparation

Type I Bovine Collagen solution (PureCol® 3 mg/ml, Advanced BIOMATRIX) was used as the bioink for cell printing. The collagen solution was neutralised following the manufacturer’s instructions to an approximate pH of 7.2, and the hiPSC cell suspension was added, resulting in a final collagen concentration of 2.4 mg/ml. The final hiPSC concentration in the bioink was 5.6 million cells/ml. Brillant Blue FCF dye (BCL3134) was added to the bioinks to enable visualisation using the computer vision system. The dye was added to the bioink at a final concentration of 0.087 mg/ml. Post-printing, the bioprinted constructs were incubated for 1h at 37°C to physically crosslink the collagen network.

### Rheological characterisation

The viscosity of the 100 mg/ml alginate ink, 50 mg/ml alginate ink, and 2.4 mg/ml collagen bioinks were measured as a function of shear rate (0 – 100 s^-1^) using rotational shear rheometry (25 mm diameter, gap 1 mm, Anton Paar MCR 302) at 25°C.

### Live/dead staining and confocal microscopy

Live/dead staining was performed on Day 0 to assess the viability of the bioprinted constructs. The bioprinted constructs were treated with 2 μM Calcein-AM (11564257 Fisher Scientific) and 4 μM Ethidium homodimer (E1903, Merck) in 1x PBS, followed by a 45 min incubation at 37°C. The stained constructs were imaged using a confocal microscope (BC43 Benchtop Confocal Microscope, ANDOR).

## Acknowledgements

This publication has emanated from research conducted with the financial support of the EU Commission Recovery and Resilience Facility under the Research Ireland Future Digital Challenge Grant Number 22/NCF/FD/10991. This publication has also emanated from research supported in part by a grant from Research Ireland and is co-funded under the European Regional Development Fund under Grant number 13/RC/2073_P2.

## Data availability

All data supporting the results of this study can be found at the following link https://doi.org/10.5281/zenodo.13939763. All code related to the print quality analysis can be found at the following link https://doi.org/10.5281/zenodo.13981630.

